# Computational prediction of CRISPR-impaired non-coding regulatory regions

**DOI:** 10.1101/2020.12.22.423923

**Authors:** Nina Baumgarten, Florian Schmidt, Martin Wegner, Marie Hebel, Manuel Kaulich, Marcel H. Schulz

## Abstract

Genome-wide CRISPR screens are becoming more widespread and allow the simultaneous interrogation of thousands of genomic regions. Although recent progress has been made in the analysis of CRISPR screens, it is still an open problem how to interpret CRISPR mutations in non-coding regions of the genome. Most of the tools concentrate on the interpretation of mutations introduced in gene coding regions. We introduce a computational pipeline that uses epigenomic information about regulatory elements for the interpretation of CRISPR mutations in non-coding regions. We illustrate our approach on the analysis of a genome-wide CRISPR screen in hTERT-RPE-1 cells and reveal novel regulatory elements that mediate chemoresistance against doxorubicin in these cells. We infer links to established and to novel chemoresistance genes. Our approach is general and can be applied on any cell type and with different CRISPR enzymes.

## Introduction

One of the largest problems in current biology and molecular medicine is understanding how genotypes cause phenotypes. The recent development of the Clustered Regularly Interspaced Short Palindromic Repeats (CRISPR) technology accelerated genotype to phenotype associations by introducing targeted gene perturbations [1]. While arrayed CRISPR experiments enabled the association of phenotypes to the perturbation of individual genes, genome-wide CRISPR experiments are able to target entire genomes simultaneously in a single experiment [2,3]. Unbiased genome-wide CRISPR screens facilitated the linking of genotypes with phenotypes on a large scale and facilitated the identification of drug resistance and cancer vulnerabilities [4–7]. In addition to fishing for potential hit candidates on a genome-wide scale, the CRISPR technology has been used to understand gene architecture and identify functional protein domains through exhaustive gene tiling [8, 9]. Here, all possible gRNAs for a locus of interest were tested simultaneously to identify gene/protein segments in which gRNAs were under/over-represented due to a phenotypic enrichment. Moreover, CRISPR screens have been applied to the non-coding genome to functionally characterize long non-coding RNAs (lncRNAs) and cis-regulatory elements linked to cell viability and chemotherapy resistance [3, 10, 11].

Current approaches to analyze genome-wide CRISPR screens are based on computing log fold changes of gRNA abundance between treated and untreated cell populations. Usually, the treated cell population is harvested at the end time point of the screen and compared to an untreated control population that has been harvested at an early time point, or even the plasmid library. Positive and negative hit gRNAs are then called based on the ranking of these relative changes [7, 12]. This principle can be applied to analyze CRISPR screens targeting protein-coding genes as well as non-coding regions. For CRISPR screens targeting known protein-coding genes, a variety of statistical analysis tools are available: One of the first algorithms that was specifically designed for the analysis of CRISPR knockout screens was MAGeCK, around which an entire suite of visualization and analysis tools has evolved [13–15]. The BAGEL algorithm aims to identify hitherto unknown essential genes by applying a statistical model that is based on prior knowledge about gene essentialities [16]. A user-friendly and easily accessible tool is PinAPL-Py which is implemented as a web service and offers multiple analysis options for a variety of CRISPR screening experiments [17]. However, there are no general approaches available to call hits of a genome-wide screen using a randomized gRNA library that targets mainly non-coding regions. The reason is that an additional annotation step is required to overlap hit gRNA target sites in non-coding regions with known or predicted gene regulatory elements. Also, it is currently unclear whether the same statistical assumptions hold true for calling hits in coding and non-coding regions of the genome.

Understanding the regulation through non-coding regions is a current topic in genetics. One particular role of the non-coding genome is to harbor Regulatory EleMents (REMs) such as enhancers, repressors, and promoters. REMs can be bound by transcription factors (TFs). TFs can recruit other proteins that can influence the 3D-structure of the chromatin and regulate gene expression in a cell-type specific manner [18]. REMs can be located far away from the genes they regulate and lead to up- or down-regulation of target gene expression [19–21]. Identifying REMs is challenging, and different genome-wide protocols can be used. There is no one gold-standard method, but instead different (epi-)genomic indicators and different algorithmic approaches are used for the annotation of REMs [22–28].

There are a number of databases that describe human REMs, such as the*Vista Enhancer Browser* [29], with experimentally verified enhancers, the FANTOM5 enhancer resource [26], or the *HACER* database [30]. Furthermore, there are resources that hold information about REMs and their putative target genes, the genes they regulate. Examples are the GeneHancer database [31] or *RAEdb* [32]. Another large-scale source of human REMs is *EpiRegio* [33], a database that contains 2.4 million predicted REMs with associated target genes predicted with the StitchIt algorithm [28]. Databases such as *HEDD* [34] or *DiseaseEnhancer* [35] list REMs that are associated to known disease genes, highlighting the role of non-coding variation in human disease.

In this study we develop and test a workflow for the prediction of non-coding CRISPR events that disrupt gene regulation based on the annotation of human REMs and epigenomic cell-specific information. We apply our approach to a genome-wide CRISPR screen that has been performed in hTERT RPE-1 cells to find coding and non-coding regions that mediate doxorubicin resistance [3].

Doxorubicin is a chemotherapy medication used to treat different forms of cancer. It induces double-strand DNA breaks and triggers DNA damage associated cell cycle arrest and apoptosis pathways, for example via MAPK/ERK pathway [36,37]. Our analysis reveals 35 genes that we could link to doxorubicin resistance through genomic pertubations in active REMs in RPE-1 cells.

## Results

### Overview of the suggested approach

Our approach identifies regulatory elements (REMs) linked to their target genes, which are (1) genomically modified by a gRNA and (2) are active in the cell type of interest. In order to identify the genomically modified REMs, we intersect them with the gRNAs binding sites. Additionally, epigenomic data measuring open chromatin, *e*.*g*. DNase1-seq, or histone modifications associated with active transcription, *e*.*g*. H3K27ac or H3K4me3, can be used to identify REMs active in the cell type of interest. Based on the active and genomically modified REMs, a protein-protein association network of the associated target genes is constructed and a motif enrichment analysis of the REMs is performed. A general overview of our approach is shown in Fig. 1.

**Fig 1.**
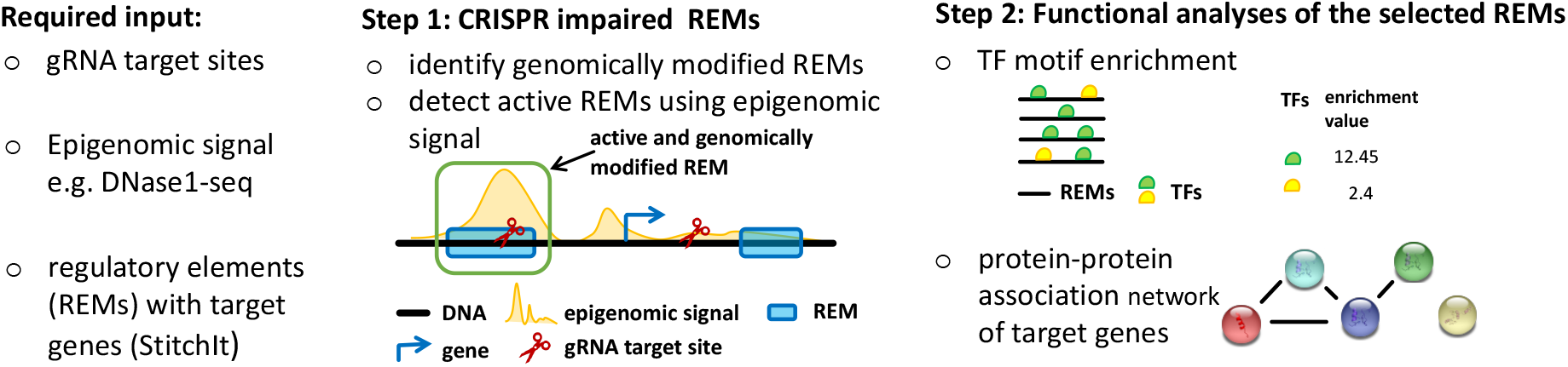
Overview of our analysis approach. Step 1: gRNA target sites are overlapped with a catalogue of gene regulatory elements (REMs) and peak regions of epigenomic measurements for regulatory activity (*e*.*g*. DNase1-seq, H3K4me3). Step 2: functional analysis of the target genes reveals involved complexes. Transcription factor (TF) motif enrichment in REMs identifies TFs that are involved in the trait of interest.

### Prediction of regulatory regions that mediate chemoresistance in RPE-1 cells

To illustrate our newly proposed approach on a realistic application, we applied it to a genome-wide doxorubicin CRISPR-Cas9 resistance screen in hTERT-RPE-1 cells by Wegner *et al*. [3]. As a result of the screen, 332 non-coding target sites of 226 validated gRNA sequences were identified. We extended the gRNA target sites to regions of length 200bp such that the target site is centered in the middle. We aim to identify REMs affected by gRNAs, and explore their possible involvement in chemoresistance.

The REMs and their target genes were inferred simultaneously by Schmidt *et al*. using StitchIt [28]. This method interprets the variation in epigenetic data, like DNase1-seq in relation to gene expression, across various cell types. To take even distal REMs into account, StitchIt ‘s predicted REMs are within a window covering 100kb upstream of the gene’s transcription start site, the gene body and 100kb downstream of the gene’s transcription termination site. In total, we downloaded 3,900,708 REMs associated to 36,817 genes based on data from the Blueprint [38], Roadmap [39] and the ENCODE [40] consortium.

We found 219 putative REMs overlapping with a non-coding gRNA, which could be linked to 190 different genes (see Sup. Table).

To ensure that we observe the overlap not only by chance, we randomly shuffled the positions of gRNA target sites within the genome and intersected the random regions with the REMs from our catalogue. We repeated this 100 times and obtained on average 72.07, 83.87 and 25.45 REMs overlapping with the sampled regions for Blueprint, Roadmap and ENCODE, StitchIt REMs, respectively. We used a two-sided *t*-test to compare results from the shuffling analyses with the REMs overlapping true gRNAs and found a significant enrichment in each data set (Blueprint: 89/72.07, *p*-value < 2.2e-16; Roadmap: 90/83.87, *p*-value 1.638e-06; ENCODE: 40/25.45, *p*-value < 2.2e-16).

### Analysis of epigenomic data reveals strong candidate doxorubicin chemoresistance genes

StitchIt identifies REMs using epigenomics data from several cell types, therefore we can not directly conclude whether or not REMs affected by a gRNA are *active* in hTERT-RPE-1 cells. To identify active REMs in open chromatin regions, we have integrated DNase1-seq and H3K4me3 ChIP-seq data from human RPE-1 cells [40]. In total, we identified 13 gRNA target sites, which are overlapping active REMs linked to 35 different genes (see Table 1). Interestingly, it occurred several times that a gRNA target site overlapped a region, which is linked to several target genes. These genes may be strong candidates to be involved in doxorubicin chemoresistance, as gRNA-mediated change of the DNA sequence may impair gene regulatory functions.

**Table 1.** Candidate doxorubicin chemoresistance genes. All gRNAs that overlap regulatory regions with epigenetic evidence and their associated target genes are listed. Genes which have been reported in the literature to be linked to chemoresistance or cancer growth are in bold.

| chr | gRNA coordinate | gene(s) associated |
| --- | --- | --- |
| 1 | 40,036,652 | CAP1, PPT1, AL663070.1 |
| 10 | 103,282,011 | **NT5C2** [41] |
| 10 | 3,197,670 | **PFKP** [42] |
| 10 | 73,358,465 | CFAP70, RPL26P6, **ANXA7** [43, 44], RNU6-833P |
| 13 | 87,671,102 | SLITRK5 |
| 16 | 30,021,011 | **TMEM219** [45], **DOC2A** [46], TBX6, **GDPD3** [46, 47] |
| 17 | 28,404,283 | AC002094.1, KRT18P55, SEBOX, **TMEM199** [48] |
| 17 | 42,423,866 | MIR5010 |
| 17 | 81,635,498 | MRPL12, **SLC25A10** [49, 50], AC137896.1, HGS, TSPAN10, AC139530.3, CCDC137 |
| 20 | 45,993,823 | NCOA5, **MMP9** [51], PCIF1 |
| 22 | 24,063,890 | AC253536.3 |
| 4 | 139,651,636 | AC112236.2, H3P16, **MGST2** [52] |
| 8 | 120,445,481 | MTBP, COL14A1 |

We have performed a detailed literature survey to find previous studies that have linked our candidate genes to chemoresistance against doxorubicin or other cancer drugs, supporting a putative role of REM target genes. We also considered studies that linked candidate genes to cancer growth, which may help to establish chemoresistance. In total, we have found that 8 out of 13 gRNAs link to at least one gene with literature evidence, as detailed below.

In myocytes, an increase of Myocardial Metallo Proteinases (MMPs) expression through increasing ROS formation induced by doxorubicin was observed [51]. At position chr20:45993823, overlap between a gRNA and REMs assume a regulatory site of MMP9, being in agreement with the finding of Spallarossa *et al*. [51]. Sun *et al*. showed that in colorectal cancer NCOA5 is upregulated, which leads to an higher expression of MMP9 and Cyclin D1 as well as to a lower expression of p27 through PI3K/AKT pathway. As a consequence, cell proliferation, migration and invasion are enhanced [53].

Another candidate is the NT5C2 gene, which is known to confer treatment resistance against thiorubin in acute lymphocytic leukemias [41]. For ANXA7 it was shown that resistance in different cancer types has been observed [43, 44].

One particularly interesting region was a gRNA binding to active REMs in a locus on chromosome 16. For three out of the four target genes we have found literature evidence. TMEM219 is a IGFBP-3 receptor that has been linked to promoting anti-tumoric effects of IGFBP-3 [45]. The gene expression and protein abundance of GDPD3 is associated with positive response to neoadjuvant chemotherapy in urothelial carcinomas [47]. A genome-wide screen for induced genes after doxorubicin in cancer cell lines [46] found that from the genes in Table 1 GDPD3 and DOC2A showed differential expression.

The expression of the TMEM199 gene was found to be upregulated in cisplatin resistent oesophageal adenocarcinoma cells pointing to a role in chemoresistance [48]. The gene product of SLC25A10 is a protein that belongs to the family of mitochondrial carriers, that are potential cancer therapy targets [49]. For example, high protein levels of SLC25A10 are associated with high metastasis potential and lower relapse-free survival in osteosarcomas [50]. Moon and others [54] found PFKP expression is upregulated in breast cancer cells. Further, high glycolytic metabolism, has been linked to breast cancer aggressiveness and growth [42].

Dvash *et al*. showed that endoplasmic reticulum stress and doxorubicin or 5-fluorouracil induce expression of MGTS2, and by that LTC4, which triggers DNA damage and cell death [52]. MGTS2-deficient mice seem to be resistance against the treatment with 5-fluorouracil. Further, they hypothesize that the MGST2-LTC4 pathway is not activated by commonly used cancer drugs, *e*.*g*. doxorubicin or 5-fluorouracil, in cells lines not able to express MGTS2, such as haematopoietic cell lines. They argue this could explain their chemoresistance.

As we found that many genes from Table 1 have been linked to chemoresistance, we wondered if they may form functional modules in the cell. We used the STRING database [55] to compute a protein-protein association network that contains all genes from Table 1 (see Fig. 2). The analysis revealed that DOC2A, TMEM219, GDPD3 and TBX6 form an association module. This is because all four genes lie within a region of 600kb, which is connected to different forms of human disease depending on whether the region is deleted or duplicated [56, 57].

**Fig 2.**
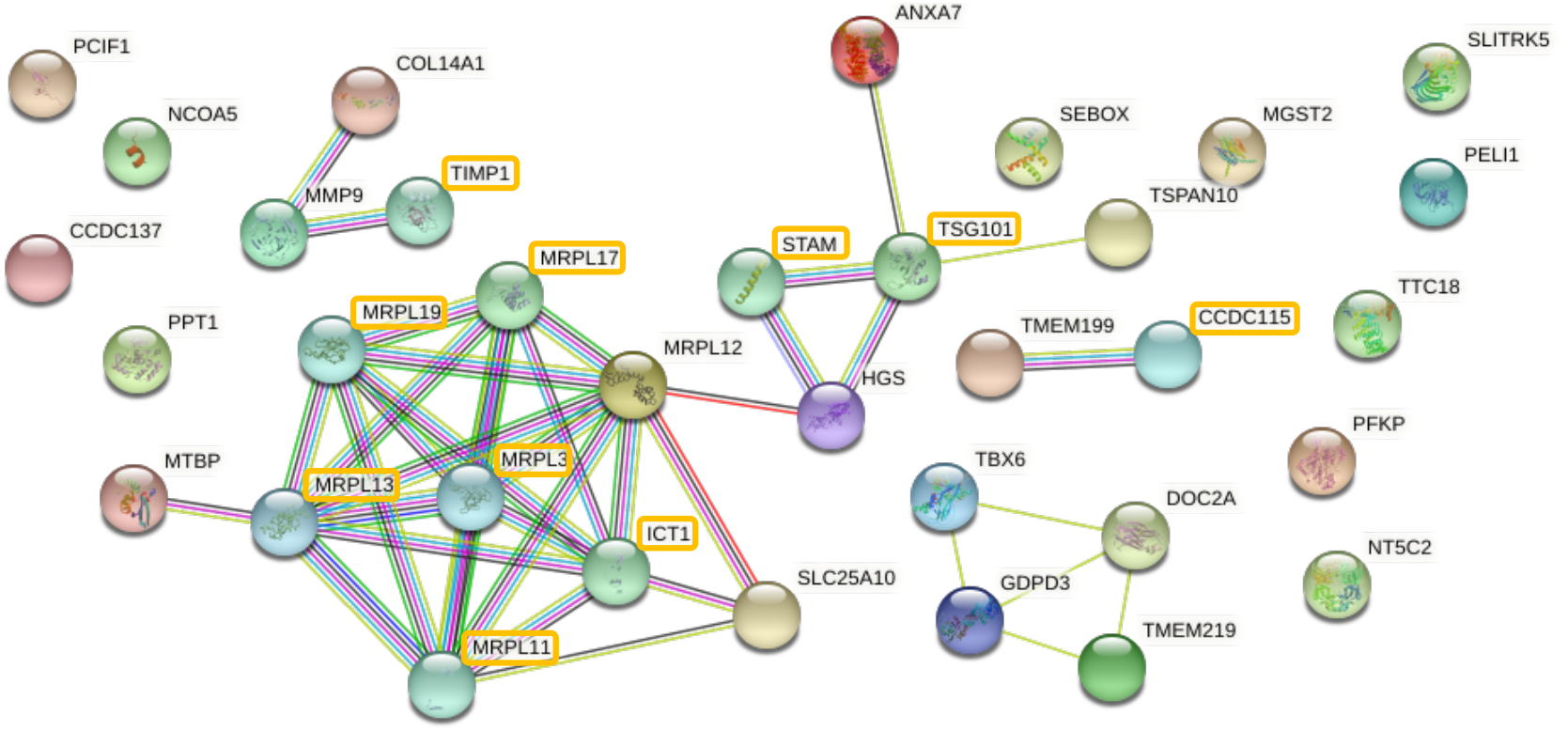
Functional analysis of target genes. Protein-protein association network of the genes associated to active, genomically modified REMs from the STRING database [55]. The network was seeded with all genes in Table 1 and includes 10 additional interactors/proteins (marked with orange box) highly connected to the genes provided as input.

### Enrichment of transcription factors binding in chemoresistance regulatory elements

To identify key TFs involved in the regulatory process of doxorubicin resistance, we performed a TF motif enrichment analysis based on the active, genomically modified REMs. The analysis is done with *PASTAA* [58], where we have ranked sequences according to the open chromatin signal to detect enriched TFs compared to background REMs (more details in the methods section).

Table 2 lists the most enriched TFs and their corresponding FDR corrected *p*-value. The table contains a number of TFs that have been linked to chemoresistance. For example NR2C2 (or testicular nuclear receptor TR4) has been shown to suppress prostate cancer invasion by preventing infiltration of macrophages [59] and a recent report showed that the sensitivity to Docetaxel chemotherapy can be improved using antagonists against NR2C2 [60] in prostate cancer. Further, the TF NR2F2 (or Chicken ovalbumin upstream promoter transcription factor II, COUP-TF2) expressed in different types of colorectal carcinoma, enhances the resistance to doxorubicin [61]. Several reviews [62–64] point to the importance nuclear receptors play during the development of cancer, *e*.*g*. breast or prostate cancer, and their potential role as drug targets. We identified several nuclear receptors within our analysis, like the before mentioned NR2F2 and NR2C2 factors, but also RARA, NR3C1, NR2C1 and THRB.

**Table 2.**
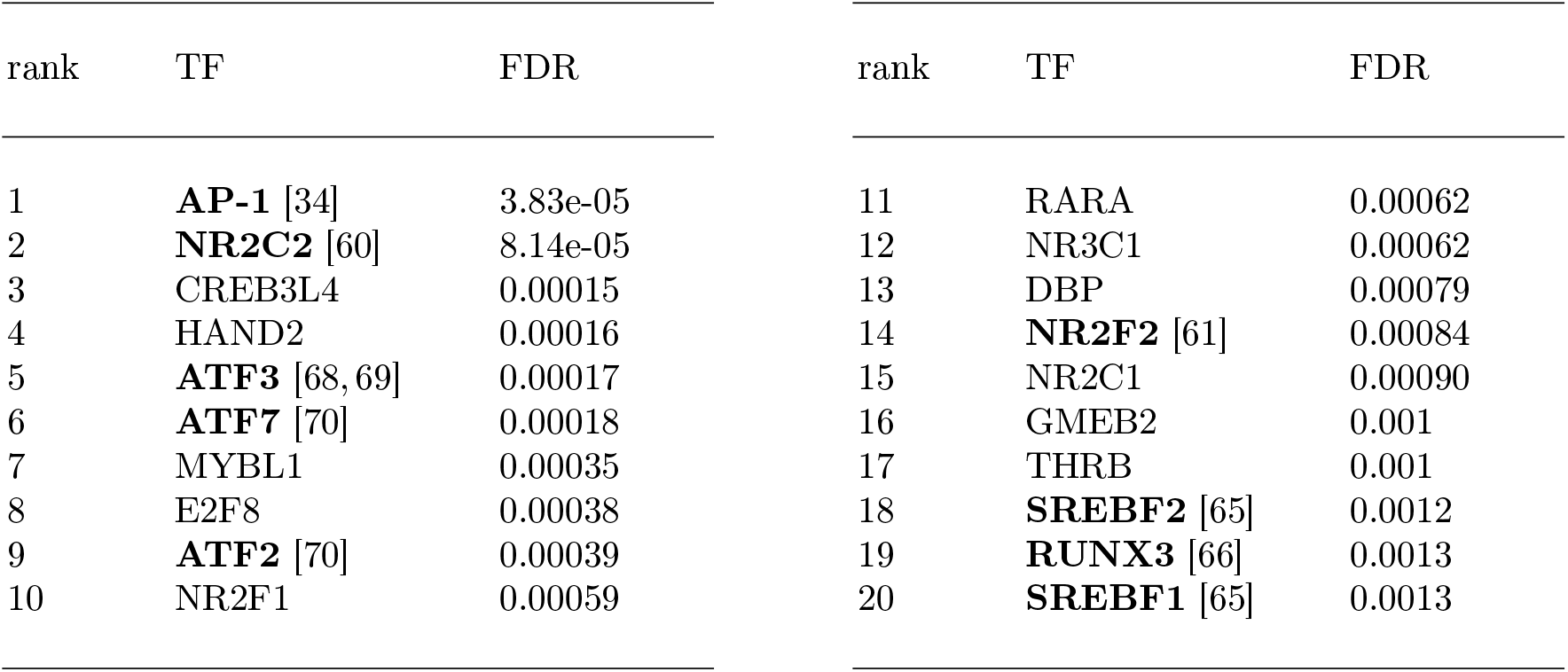
Results of the TF enrichment analysis. TF binding in genomically modified active REMs was contrasted to random sets of active REMs in the same cells. Reported are the ranked TF names and FDR corrected enrichment *p*-values. TFs which have been reported to be linked to chemoresistance are in bold.

A recent study [65] identified the G protein-coupled receptor 120 (GPR120) as a chemoresistance-promoting factor in breast cancer. The authors hypothesize that GPR120 mediates fatty acid synthesis and identified several factors, like SREBF1 and SREBF2, involved in this process. Due to the increased lipid synthesis this leads to a different lipid composition of the cell membrane and may prevent the drug from being absorbed.

Zheng *et al*. showed that the down regulation of RUNX3 in human lung adenocarcinoma is associated with docetaxel resistance by activating Akt1-mediated signaling [66]. Over-expression of RUNX3 leads to low AKT expression and higher treatment sensitivity.

ATF3 is a TF known to be involved in cellular stress response and is enriched in cells exposed to stress signals [67]. Under doxorubicin treatment, it has been reported that ATF3 affects cell death and cell cycle progression, however it is unclear whether the factor acts as a negative or positive regulator [68,69]. Nobori *et al*. claim that ATF3 plays a pivotal role as transcriptional regulator in the process of doxorubicin-induced cytotoxicity via an ERK-dependent pathway.

Walczynski *et al*. showed that the loss of ATF2 and ATF7, known to be homologous members in the AP-1 (activator protein 1) family, in MYC-expressing lymphoma cells in a mouse model are more resistant to doxorubicin induced apoptosis than control cells [70]. Further, they claim that ATF2/7 are important key TFs to suppress an oncogenic transformation. It has been found in other studies that upregulation of AP-1 expression confers resistance to chemotherapeutics [34].

## Discussion

We proposed a approach to identify genes whose expression might be affected by gRNAs targeting non-coding regulatory regions. To do so, we examined which gRNAs modify regulatory elements associated to potential target genes. The regulatory elements were retrieved from StitchIt. Further, we took epigenetic data into account to identify REMs in open-chromatin or active enhancer regions of RPE-1 cells. Based on the target genes of active and genomically modified REMs, we constructed a protein-protein association network. Using the REM sequences, we performed a TF motif enrichment analysis to obtain key TFs involved in regulating doxorubicin resistance (see Fig. 1). Several of the identified target genes and TFs are already known to be associated to doxorubicin resistance, chemoresistance or cancer growth in general (marked in bold in Table 1 and Table 2).

Nevertheless, there are genes and TFs remaining that are worth to investigate more closely. For instance, the TF motif enrichment analysis points to the factor CREB3L4, for which we were not able to find an association to doxorubicin resistance. However, protein abundance of CREB3L1, which belongs to the same TF subfamily as CREB3L4, is predictive of the response of triple negative breast cancer to doxorubicin-based therapy [71]. In a recent paper, Pu *et al*. already identified that expression of CREB4L3 is increased in most breast cancer types, and that the downregulation leads to a decreasing proliferation, induces cell cycle arrest and apoptosis [72]. We observed a CREB3L4 binding site in a genomically modified REM associated to CCDC137. The gRNA (chr17:81,635,498) also affects MRPL12, SLC25A10, AC137896.1, HGS, TSPAN10 and AC139530.3.

We found one example, where a gRNA target site overlaps with the REM of the post-transcriptional regulator gene MIR5010 (Table 1). Mature microRNAs produced from this locus have the capacity to regulate many target genes. While we were not able to find evidence that the mature microRNA products hsa-miR-5010-3p or hsa-miR-5010-5p of the MIR5010 gene are linked to chemoresistance against doxorubicin, the prevalence of hsa-miR-5010-3p may be involved in colon cancer recurrence. In a study of TNM-stage II colon cancer patients in two different cohorts, a four-miRNA recurrence prognosis signature was developed, including the abundance of hsa-miR-5010-3p [73]. Other than that, little is known about the regulatory roles of mature microRNAs of MIR5010. However, we queried the SpongeDB database, that holds predicted competing endogenous RNA interactions computed for different cancer types [74] using the Sponge algorithm [75]. We found that hsa-miR-5010-3p was predicted to regulate 3455 gene-gene ceRNA pairs in a pancancer model and many others in individual cancer types (Suppl. Tab. 1), suggesting an important regulatory role in cancer.

The overall association approach introduced here differs from other ad-hoc approaches. For example Wegner *et al*. identified target genes using the nearest gene approach, which links non-coding regions to adjacent genes without additional evidence. Our newly proposed approach is based on epigenetic data, first to infer the regulatory regions and second to detect genomically modified REMs, which are accessible in RPE-1 cells. Therefore, we conclude that our strategy is more informative than an approach, which is just based on genomic locations.

A CRISPR modification can create new TF binding sites in REMs. In rare cases, a consequence is that inactive REMs in the cell type of interest can become active by the gRNA caused modification. In general, no open chromatin data is available for the cell types after the modification. For that our current approach can not identify the REMs which become active by gaining a TF binding site.

We are aware that a CRISPR-Cas9 experiment can lead to off-target effects. Since we were able to link suggested target genes and TFs to chemoresistance events, we assume that at least for the considered cases here, off-target effects do not play a major role. Following the assumption that no strong off-target effects are causing chemoresistance in the experiment, it would be remarkable that only the modification of one REM leads to chemoresistance. This may fit to the observation that many of the active REMs detected here are likely regulating more than one target gene.

While we wanted to use ENCODE epigenomic data for RPE-1 cells, predictions of StitchIt and additional open-chromatin and expression data for the Roadmap and Blueprint consortium can be directly obtained from our *EpiRegio* webserver [33]. *EpiRegio* holds data for diverse cell types, which allows to retrieve active REMs with their associated target genes.

To sum up, we introduced a cell-type specific approach to identify potential target genes of CRISPR modifications in non-coding regions, and useful downstream applications such as TF motif enrichment analysis. Several of our identified target genes and TFs are known to be associated to doxorubicin resistance, chemoresistance and cell growth in cancer cells. We believe that our approach based on epigenetic data can be helpful to identify novel target genes for different CRISPR screens and various cell types.

## Materials and methods

### Analysis of Doxorubicin resistance in hTERT-RPE1 cells using CRISPR experiments

Here, we use a genome-wide CRISPR perturbation library consisting of partially randomized degenerated oligonucleotides (5’-NNDNNNNNHNNNNHDHNVVR-3’) with flanking 3Cs homology regions which was created using ssDNA of template-plasmids and site-specific mutagenesis targeting coding and non-coding regions of the human genome in hTERT-RPE1 cells from ATCC (CRL-4000) [3]. Genomic coordiantes of the gRNAs have been obtained from this work (see Supp. Tab. 1).

### Retrieval of epigenomic data

StitchIt was applied to three data sets derived from different consortia, namely Roadmap [39], Blueprint and ENCODE [40]. We downloaded the REMs predicted for the 100kb window from ZENODO (DOI: 10.5281/zenodo.4316356).

Narrow peak files for human RPE-1 cells of a DNase1-seq and a H3K4me3 ChIP-seq data set were retrieved from ENCODE (accession numbers: ENCFF535GNR, ENCFF749ZBH, ENCFF922ERE). For the DNase1-seq the data contained 140,038 and 118,815 peaks for replicate 1 and 2, respectively. The replicates show a high agreement. The H3K4me3 ChIP-seq data included 34,179 peaks.

### Identification of active and genomically modified REMs

First, we extended the gRNA target sites (see Sup. Table) to regions of length 200bp by adding 100bp up- and downstream. To identify the REMs which are active, and genomically modified by the extended gRNAs, we applied the following bedtools commands:

`bedtools intersect -a <REMs> -b <gRNAs> -wa -u > <outputFile>`

`bedtools intersect -a <outputFile> -b <H3K4me2-peaks> -wo`

`bedtools intersect -a <outputFile> -b <DNase1-seq-rep1>, <DNase1-seq-rep1> -wo`

### Details about the motif enrichment analysis

The motif enrichment analysis is performed with *PASTAA* [58], which requires the DNA sequences of the 45 REMs and a set of known TF binding motifs. We determined the DNA sequences using the *getfasta* functionality in bedtools [76] and retrieved 515 TF binding motifs from the JASPAR motif database [77]. Additionally, *PASTAA* requests a ranking of the DNA sequences. Hence, we sorted the sequences in descending order based on their maximal epigenetic signal from either the DNase1-seq or the H3K4me3 ChIP-seq, such that regions with higher read coverage occurred at the top. To ensure that PASTAA is able to discriminate enriched TF binding sites, we added 3 times as many randomly selected REMs (135) not affected by a gRNA, but with epigenomic evidence as being active in RPE1-cells. The background sequences are sorted in ascending order according to their epigenomic signal, and added to the bottom of the ranking the active, genomically modified REMs. We repeated the motif enrichment analysis 100 times, each run with a different background set, and averaged the TF enrichment results. The averaged result can be found in the Sup. Table.

## Acknowledgements

We are thankful to the ENCODE consortia for sharing the epigenomic data used in this work.

